# siRNAs targeting a chromatin-associated RNA induce its transcriptional silencing in human cells

**DOI:** 10.1101/2022.05.11.491477

**Authors:** Julien Ouvrard, Lisa Muniz, Estelle Nicolas, Didier Trouche

**Author notes:** Correspondence to : Didier Trouche, MCD UMR 5077, Centre de Biologie Intégrative, Université Paul Sabatier, 118 Route de Narbonne, 31062 Toulouse Cedex, France. These authors contributed equally to this work (Estelle Nicolas, Didier Trouche).

## Abstract

Transcriptional gene silencing by small interfering RNAs (siRNAs) has been widely described in various species, such as plants or fission yeasts. In mammals, its extent remained somewhat debated. Previous studies showed that siRNAs targeting gene promoters can induce the silencing of the targeted promoter, although the involvement of off-target mechanisms was also suggested. Here, by nascent RNA capture and RNA Pol II ChIP, we show that siRNAs targeting a chromatin-associated non-coding RNA induce its transcriptional silencing. Deletion of the sequence targeted by one of these siRNAs on the two alleles by genome editing further show that this silencing is due to base pairing of the siRNA to the target. Moreover, by using cells with heterozygous deletion of the target sequence, we show that only the wild type allele, but not the deleted allele, is silenced by the siRNA, indicating that transcriptional silencing occurs only in cis. Finally, we demonstrate that both Ago1 and Ago2 are involved in this transcriptional silencing. Altogether, our data demonstrate that siRNAs targeting a chromatin-associated RNA at distance from its promoter induce its transcriptional silencing. Our results thus extend the possible repertoire of endogenous or exogenous interfering RNAs.

## Introduction

RNA interference (RNAi), first discovered in plants during the 1990s and observed later in other systems, like worms, yeast and mammals is a rapid and efficient tool to induce the knockdown of specific transcripts. This main mechanism of post-transcriptional gene silencing is largely conserved throughout evolution. In mammals and non-mammalian organisms, exogenous or endogenously synthetized small double-stranded RNA molecules of 19 to 24 nucleotides, called small-interfering RNA (siRNA), are loaded on the RISC (RNA-induced silencing complex) complex containing argonaute proteins (Ago) (1). The two strands of siRNAs are then dissociated and the passenger strand is ejected. The complex is then tethered to target RNAs containing a sequence complementary to the guide strand. When sequence complementarity is high enough, the endoribonuclease activity present in Ago2 cleaves the target RNA between bases 10 and 11 of the siRNA complementary region, leading to the degradation of the target RNA (2, 3).

Given that the above-described processes usually take place in the cytoplasm, RNA interference is in this respect a cytoplasmic process functioning at a post-transcriptional level, and leading to RNA degradation. Nevertheless, in some organisms such as yeast or plants, RNAi has been described to work also in the nucleus to silence repeated elements and to nucleate heterochromatin (4). However, the presence of the nuclear RNAi in mammals is more controversial. Indeed, initial data concluded that intronic sequences cannot be targeted with RNAi (5, 6). This observation together with the findings that some RISC proteins were mostly cytoplasmic led to the conclusion that RNA interference is restricted to the cytoplasm in human cells (5). However, in the last two decades, various studies have suggested the presence and the action of the RNAi machinery in the nucleus of mammalian cells. Indeed, whereas siRNA loading factors into RISC are exclusively cytoplasmic, other RNAi factors, such as Dicer, Ago1 and Ago2, have been found in the nucleus and can lead to the siRNA-mediated cleavage of chromatin-associated RNAs (7–9). Moreover, the preferential nuclear localization of Ago2 has been evidenced in specific human tissues (10). It was later found that Ago-miRNA complexes can target and silence post-transcriptionally thousands of pre-mRNAs in the nucleus in mouse ES cells that contain high levels of nuclear Ago proteins (11, 12), although it is still unclear whether all these genes are directly targeted by Ago proteins. Co-transcriptional processes such as alternative splicing can also be affected by siRNA targeting a specific splice site of pre-mRNA [39] Moreover, siRNAs have been successfully used to decrease the expression of nuclear ncRNAs (7, 13), including by ourselves (14–16).

In addition, transcriptional gene silencing by siRNAs was also reported in mammalian cells. siRNAs directed against the promoter of the elongation factor 1α gene repress the expression of a reporter gene driven by this promoter (17). Further studies confirmed for other genes that siRNAs can repress transcription when targeting promoters (18–25) or first exons (13). This was also extended to miRNAs or siRNAs targeting downstream regions of protein-coding genes (26, 27). Transcriptional gene silencing requires an RNA template (28, 29), such as promoter-associated antisense transcripts (30) or ncRNAs overlapping the gene promoter or downstream regions (25, 31) and involves either Ago1 or Ago2 (18, 31, 32). The targeting of the siRNA to promoters leads to the deposition of repressive chromatin marks at these promoters, such as H3K9me3 and H3K27me3 (9, 20, 24, 32–34) or CpG methylation (19, 22), resulting in transcriptional gene repression. Note that it was proposed that the previously reported transcriptional repression by siRNAs of the VEGF gene promoter and HIV LTR could be due to off-target effects, since a similar effect was observed on reporter systems with a deleted or mutated target site (35). For the HIV LTR, a later study showed that siRNAs targeting HIV1 LTR are specific to the HIV1 LTR and do not affect the highly related HIV2 LTR neither a panel of other genes regulated in a similar way, arguing against purely off target effects (36).

Despite these studies, to our knowledge, siRNAs repressing promoters while targeting RNAs far downstream the promoter region has not been reported so far. Here, we show that siRNAs directed against a chromatin-associated RNA from the vlincRNA (very long intergenic non-coding RNA) family can silence its promoter even when targeting sequences are located thousands of bases downstream from the promoter. Moreover, through the use of genome editing, we found that this effect occurs in cis and is not due to off target mechanisms, therefore demonstrating RNA interference mechanisms acting at distance in mammals.

## Results

### siRNAs can efficiently decrease the expression of VINK nuclear RNA

In previous works, we successfully used siRNAs to deplete the expression of various RNAs associated with chromatin (14–16). This includes depletion of VAD, a senescence-specific ncRNA which belongs to the vlincRNA family (16). More recently, we investigated another vlincRNA also strongly associated with chromatin with more than 85 % present in the chromatin fraction and very weakly spliced, which we named VINK (Ouvrard et al., in preparation). During the course of this investigation, we designed and transfected siRNAs directed against this vlincRNA in RAF1 oncogene-induced senescent WI38 human cells. By RT-qPCR, we observed that an siRNA against VINK (VINK-1 siRNA) efficiently inhibits VINK expression when measured at various places along the VINK RNA (Figure 1A and 1B). A similar result was obtained with a second independent couple of control and VINK siRNAs (VINK-2 siRNA), although VINK inhibition was slightly less efficient with this siRNA (Figure 1A and 1C). A dose response showed that the effect was maximal from 10 nM of siRNA (data not shown). The efficient depletion of VINK using each of these two siRNAs was also observed on RNA-Seq experiments (Figure 1D). Thus, siRNAs against the VINK vlincRNA efficiently repress VINK expression.

**Figure 1.**
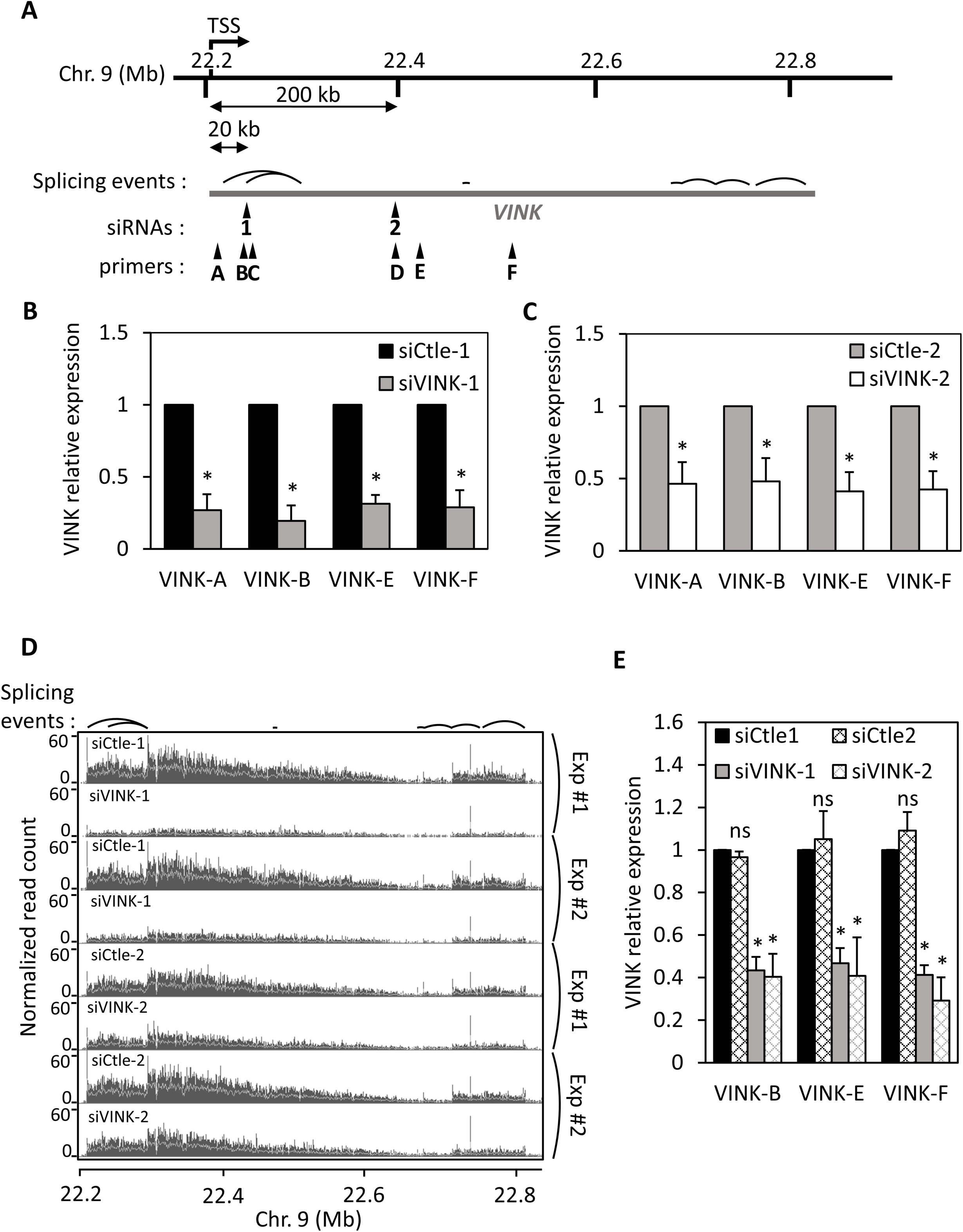
Small interfering RNAs can efficiently decrease VINK expression. **A**: Schematic representation of VINK genomic locus, with the localization of siRNA binding sites, qPCR primers, and main splicing events detected in Fig. 1D RNA-Seq experiments indicated. The exact locations of siRNAs, primers and main splicing events relative to the TSS of VINK estimated from RNA-Seq read coverage (Ouvrard et al., manuscript in preparation) are indicated in Table S1, S2 and S3, respectively. **B**: Senescent WI38-RAF1 cells were transfected with the VINK-1 or control (Ctle-1) siRNAs. 72 hours later, total RNA was prepared and analysed by RT-qPCR for the expression of GAPDH mRNA or VINK (using the indicated primers). The amount of VINK RNA was standardized to GAPDH mRNA and calculated relative to 1 for cells transfected with the Ctle-1 siRNA. The means and standard deviations from 5 independent experiments are shown. *: p value < 0.05 compared to control. **C**: Same as in B, except that siVINK-2 and another control siRNA (Ctle-2) were used. The means and standard deviations from 7 independent experiments are shown. *: p value < 0.05 compared to control. **D**: WI38-RAF1 cells were treated as in B and C, and total RNA was analysed by RNA-Seq. Two duplicates of RNA-seq experiments were visualized in the IGB browser showing the normalized read coverage for the VINK locus. Splicing events consistently detected in RNA-Seq experiments are indicated above the profiles. **E**: U2OS cells were transfected with the indicated siRNAs. 48 hours later, total RNA was prepared and analysed by RT-qPCR for the expression of GAPDH mRNA or VINK (using the indicated primers). The amount of VINK RNA was standardised to GAPDH mRNA and calculated relative to 1 for cells transfected with the Ctle-1 siRNA. The means and standard deviations from 3 independent experiments are shown. *: p value < 0.05 compared to Ctle-1 siRNA ; ns: non-significant, p value > 0.1.

From these RNA-seq datasets, analysis of reads containing at least one spliced junction showed that VINK is indeed poorly spliced with less than 0.4 % of spliced reads relative to all reads aligned in VINK region. For comparison, we previously showed that the median percentage of spliced reads in annotated genes is around 20 % (15). The main splicing junctions are indicated in Fig1A and listed in Table S3 along with the locations of primers and siRNA sequences. The most abundant splicing event occurs between a region close to the TSS and a region around 100 kb downstream (see the sharp peaks in RNA –seq datasets in Fig. 1D and Table S3).

Data showing that siRNAs can efficiently decrease VINK expression, as well as the expression of other nuclear RNAs, which we have published previously (14–16), were obtained in senescent WI38 cells expressing the hTERT, the catalytic subunit of the human telomerase and an ER-RAF1 fusion protein. To test whether the depletion of VINK RNA by siRNAs can be observed in other cell types, we made use of the human osteosarcoma U2OS cell line that also expresses VINK. In U2OS cells, VINK is truncated from its 5’end, the transcript starting around 75 nt upstream the region targeted by the siRNA-1 (data non shown). Transfection of each of the two siRNAs against VINK efficiently repressed VINK expression (Figure 1E).

Thus altogether, these data indicate that siRNAs can efficiently induce the knock-down of the chromatin-associated VINK RNA

### siRNAs affect the transcription of their targeted nuclear RNAs

Given that the use of siRNAs to target nuclear RNAs is debated, we intended to investigate the mechanism of this effect. To that goal, we first analyzed whether siRNAs directed against VINK induced its degradation. We transfected senescent WI38 cells with the VINK-1 siRNA (which is the most efficient, see Figure 1B) or control siRNA and treated transfected cells with actinomycin D to block the transcription. The decrease of VINK expression along time reflects its stability. Using this experimental setting, we found that the half-life of VINK was between 100 and 200 minutes, depending on the region of VINK analyzed (Figure 2A). This half-life was longer when we analyzed the 5’ extremity of VINK (VINK-A and VINK-B), certainly because these regions are found in spliced forms of VINK (Fig 1A Suppl Tables). However, wherever investigated, VINK half-life was not significantly changed in cells transfected with the VINK-1 siRNA. Moreover, no difference in its half-life was observed upon VINK-1 siRNA transfection following either short or long periods of actinomycin treatment (Figure 2A and data not shown). Thus, these experiments suggest that the stability of VINK RNA is not affected by the presence of VINK siRNAs.

**Figure 2.**
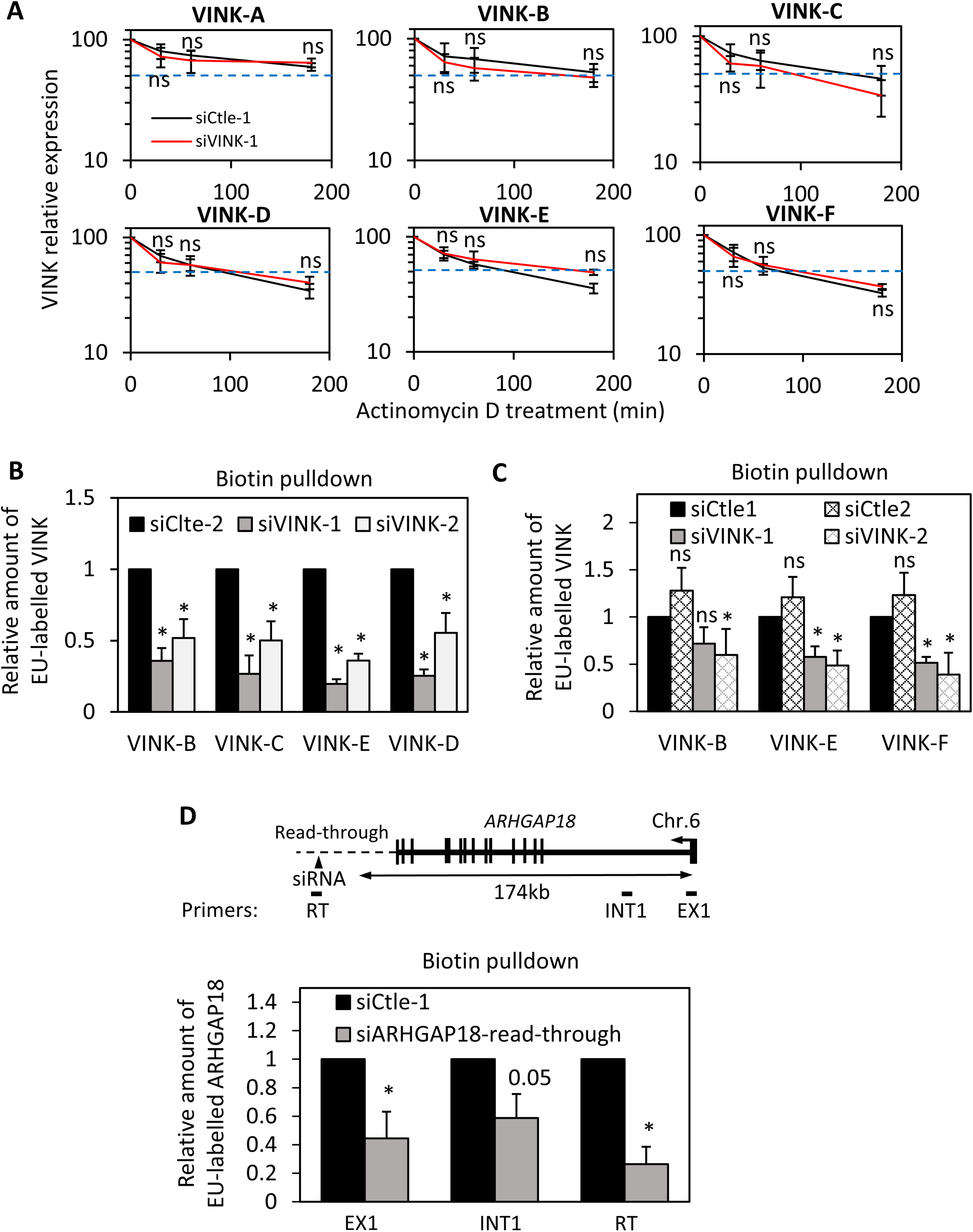
Small interfering RNAs affect VINK transcription. **A**: Senescent WI38-RAF1 cells were transfected with the VINK-1 or control (Ctle-1) siRNAs. 48 hours later, actinomycin D (10 µg/mL) was added or not. Cells were harvested after the indicated time. Total RNA was prepared with Trizol and analysed by RT-qPCR for the expression of GAPDH mRNA or VINK (using the indicated primers). The amount of VINK RNA was standardised to GAPDH mRNA (which does not vary during this time course of actinomycin D) and calculated relative to 100 for the untreated sample. The means and standard deviations from 3 independent experiments are shown. Student t test was performed to compare VINK1 or Ctle-1 siRNAs samples at each time point of actinomycin D treatment. ns: p value > 0.1. **B**: Senescent WI38-RAF1 cells were transfected with the indicated siRNAs. Nascent RNAs (Biotin-pull down) were prepared and analysed by RT-qPCR for the expression of GAPDH mRNA or VINK (using the indicated primers). The amount of VINK RNA was standardised to GAPDH mRNA and calculated relative to 1 for the Ctle-1 siRNA. The means and standard deviations from 3 independent experiments are shown. *: p value < 0.05 compared to Ctle-2 siRNA. **C**: Same as in B, except that U2OS cells were used. *: p value < 0.05 compared to Ctle-1 siRNA ; ns: non-significant, p value > 0.1. **D**: Senescent WI38-RAF1 cells were transfected with Ctle-1 siRNA or an siRNA directed against ARHGAP18 START RNA (14). Nascent RNAs (Biotin-pull down) were prepared and analysed by RT-qPCR for the expression of GAPDH mRNA or the indicated primers on ARHGAP18. The amount of ARHGAP18 or START RNA was standardised to GAPDH mRNA and calculated relative to 1 for the Ctle-1 siRNA. The means and standard deviations from 3 independent experiments are shown. *: p value < 0.05 compared to Ctle-1 siRNA ; ns: non-significant, p value > 0.1; number: p value compared to Ctle-1 siRNA.

We next assessed whether VINK siRNAs affected VINK transcription. To that goal, we analysed ongoing transcription by nascent RNA capture, which relies on metabolic labelling of RNAs being transcribed with a nucleotide analogue, allowing their biotinylation and purification on streptavidin beads. Analysis of recovered nascent RNAs thus provides an accurate measure of ongoing transcription. We transfected senescent WI38 cells with each of the two siRNAs against VINK and measured nascent transcripts at different locations along VINK (Figure 2B). We found that the presence of VINK nascent RNA was strongly decreased upon transfection of the two VINK siRNAs in all regions analyzed (Figure 2B). Thus, siRNAs targeting VINK RNA decrease its transcription. This effect was also observed in U2OS cells (Figure 2C), indicating that it is not a specific feature of the WI38 cell line.

To test whether this effect is restricted to VINK or to RNAs from the vlincRNA family, we transfected senescent WI38 cells with a siRNA directed against the ARHGAP18 read-through RNA belonging to the family of START RNAs that we characterized in a previous study (14). Read-through RNAs are produced by transcription beyond the transcription termination site and are known to be nuclear/chromatin associated (37). We also observed that the presence of ARHGAP18 START nascent RNA was strongly decreased upon transfection of the siRNA targeting it (Figure 2D). ARHGAP18 nascent transcript from which the START RNA originates was also decreased, as measured close to its TSS in its first exon and intron located about 170 kb upstream the siRNA targeted site (Figure 2D). These results indicate that the effect of the siRNA directed against ARHGAP18 START RNA is also transcriptional. Thus, we conclude from these experiments that siRNAs can repress the transcription of the chromatin-associated RNA they target, even when the targeted site is far from the promoter.

### siRNAs targeting VINK RNA influence RNA polymerase II presence at VINK TSS

Inhibition of nascent RNA upon siVINK transfection was observed throughout the VINK transcribed region, including when analyzing regions upstream of the siRNA-targeted sequence (Figures 1 and 2). siRNA-mediated inhibition of nascent transcription by premature transcription termination was recently observed in drosophila (38). To analyse whether VINK siRNAs induce premature transcription termination or involve the repression of VINK promoter, we analyzed the recruitment of RNA pol II to TSS which would be unaffected by the induction of premature termination. We thus transfected the VINK-1 or VINK-2 siRNAs and performed chromatin immunoprecipitation (ChIP) using an antibody that recognizes total RNA pol II. We found that both siRNAs against VINK inhibited the presence of RNA pol II at VINK TSS compared to control GAPDH TSS (Figure 3B and data not shown for VINK-2 siRNA), indicating that the VINK siRNAs affect RNA pol II recruitment at VINK promoter. We next analyzed whether this was also associated with the regulation of chromatin marks. We found that the VINK-1 siRNA decreased the presence of the transcription-associated H3K27ac, H3K4me3 and H3K36me3 marks at VINK TSS, whereas the repressive mark H3K9me3 was unaffected (Figure 3C). H3K27me3 was undetectable at VINK TSS, even in the presence of VINK-1 siRNA (data not shown). Thus, these data indicate that siRNAs targeting VINK decrease the presence of transcription-associated chromatin marks and of RNA Pol II at VINK TSS, therefore decreasing VINK transcription. To our knowledge, such an effect on a promoter chromatin landscape induced by siRNAs targeting RNAs thousands of bases downstream has never been reported so far.

**Figure 3.**
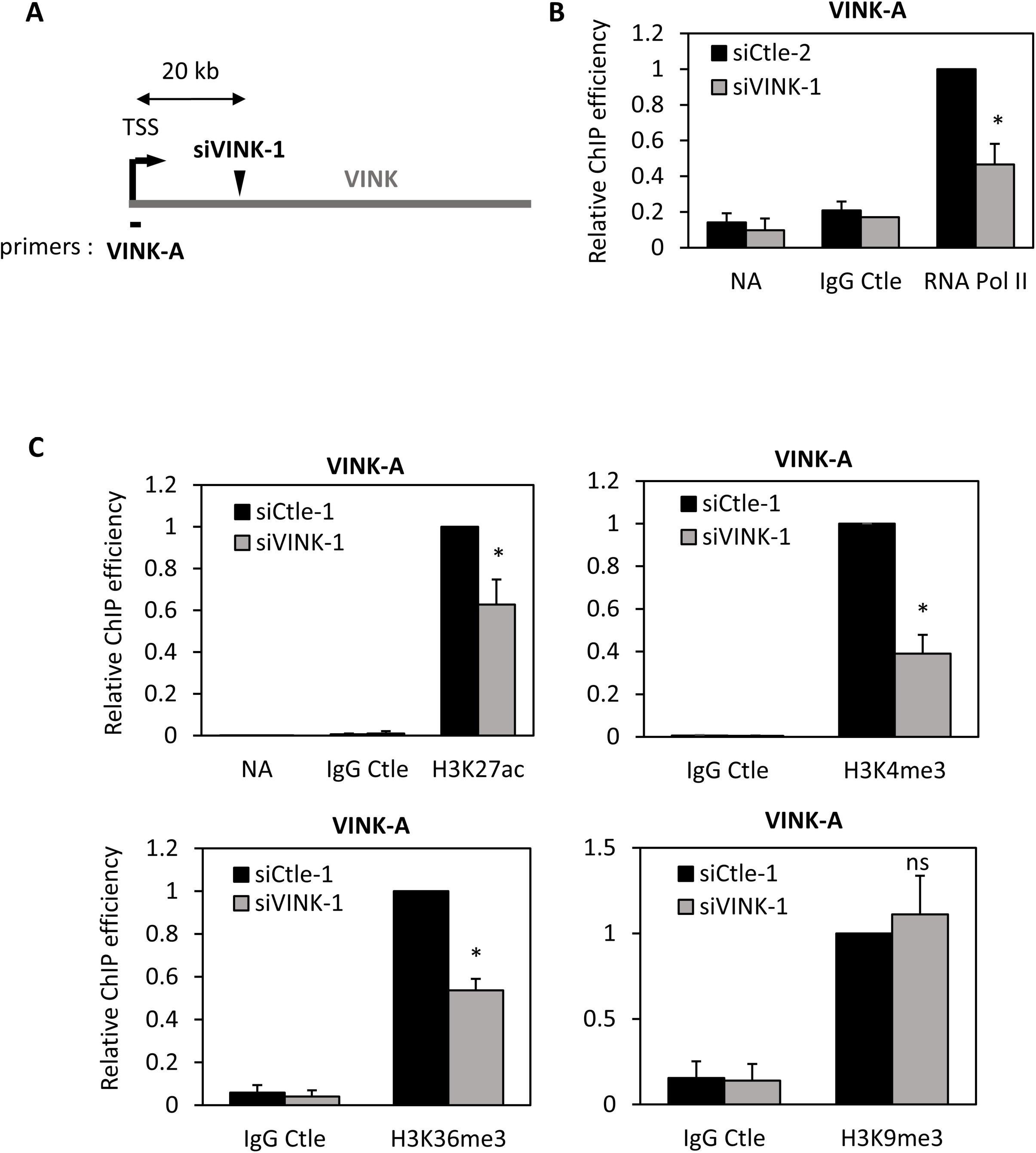
VINK siRNAs affect the presence of RNA polymerase II at VINK TSS. **A**: Schematic representation showing the localization of the TSS (transcriptional start site) of VINK and primers used to analyse ChIP experiments. **B**: Senescent WI38-RAF1 cells were transfected with the indicated siRNAs (four experiments were performed with Ctle-2 and two with Ctle-1). 72 hours later, fixed chromatin was prepared and subjected to a ChIP experiment using an antibody directed against total RNA pol II or control IgG or no antibody as a control, as indicated. The amount of VINK TSS sequence was measured by qPCR with the indicated primers, standardized to the amount of GAPDH TSS sequence and calculated relative to 1 for the RNA pol II ChIP from cells transfected by the Ctle siRNA. The means and standard deviations from six independent experiments are shown (four controlled with no antibody (NoAb) and two with a control IgG. **C**: Same as in B, except that anti-H3K27ac, anti-H3K4me3, anti-H3K36me3, anti-H3K9me3 antibodies were used. The Ctle siRNAs were the siCtle-1 siRNA except for H3K27ac experiments for which three experiments were performed with Ctle-1 and three with Ctle-2. The amount of VINK TSS sequence was standardised to the amount of GAPDH TSS sequence and to nucleosome occupancy measured by ChIP using an antibody recognising total histone H3. The result was calculated relative to 1 for relevant ChIPs from cells transfected by the Ctle siRNA. The means and standard deviations from six (H3K27ac) or three (all other antibodies) independent experiments are shown. *: p value < 0.05 compared to Ctle siRNA.

### Repression of transcription by VINK siRNA is dependent on base-pairing to its target site

We next tested whether the effect of VINK siRNA on VINK transcription is due to base-pairing and not to off-target effects. Indeed, in some instances, transcriptional silencing using siRNAs was attributed to off-target effects (35).

To analyze whether pairing of the siRNA to its target site is involved, we raised, by genome editing in WI38 cells, a stable cell line that we called WI38-Δ/Δ #1, in which 210 bp encompassing the target site for the VINK-1 siRNA are deleted on the two alleles (data not shown). We next transfected WI38-Δ/Δ #1 cells with the siRNAs directed against *VINK*. Whereas the VINK-2 siRNA works as in parental cells (compare Figures 4A and 1C), the VINK-1 siRNA, for which the sequence is not present in WI38-Δ/Δ #1 cells, did not repress VINK expression (Figure 4A). Importantly, a similar result was also observed in another independent clone deleted for the VINK-1-targeted sequence on the two alleles (WI38-Δ/Δ #2) (Figure 4B). The absence of effects of the VINK-1 siRNA on VINK expression in WI38-Δ/Δ #2 cells was also obvious from RNA-seq data when comparing the effect of the VINK-1 siRNA in parental WI38 (+/+) and deleted WI38-Δ/Δ #2 cells (Figure 4C).

**Figure 4.**
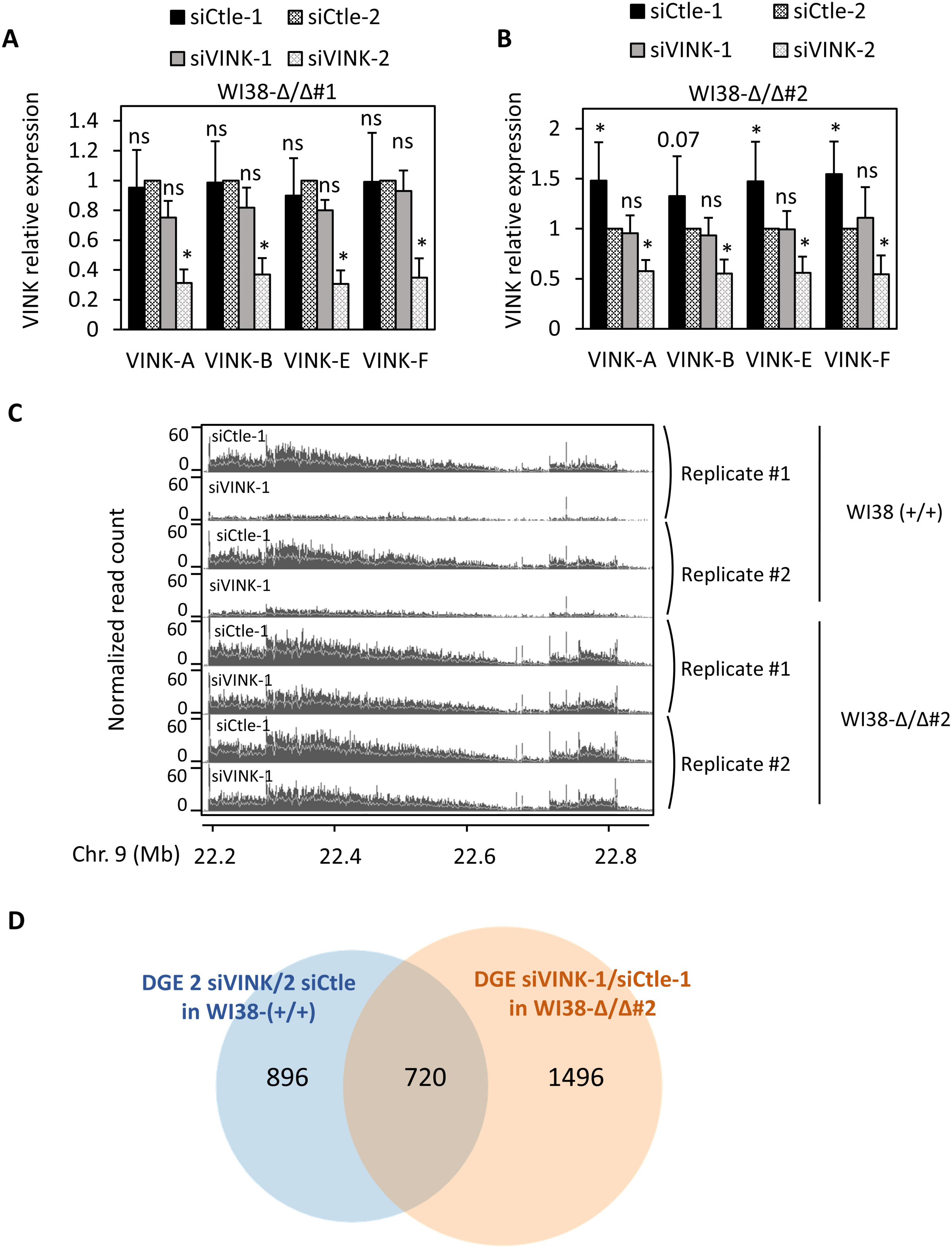
The siRNA target site is required for VINK repression. **A**: Senescent WI38-Δ/Δ #1 cells were transfected with the indicated siRNAs. 72 hours later, total RNA was prepared and analysed by RT-qPCR for the expression of GAPDH mRNA or VINK (using the indicated primers). The amount of VINK RNA was standardized to GAPDH mRNA and calculated relative to 1 for cells transfected with the Ctle-1 siRNA. The means and standard deviations from 3 independent experiments are shown. *: p value < 0.05 compared to Ctle-2 siRNA ; ns: non-significant, p value > 0.1. **B**: Same as in A, except that senescent WI38-Δ/Δ #2 cells were used. The means and standard deviations from five independent experiments are shown. *: p value < 0.05 compared to Ctle-2 siRNA ; ns: non-significant, p value > 0.1; number: p value compared to Ctle-2 siRNA. Note that siCtle1 leads to a slightly higher VINK expression than the siCtle2 in the cell line, probably due to off target effects exemplified in D. **C**: Senescent parental WI38-RAF1 (WI38-RAF1 WT) or WI38-Δ/Δ #2 cells were transfected with the indicated siRNA, and total RNA was analysed by RNA-Seq. Two duplicates of RNA-seq data for the VINK locus are shown. Note that data from WI38-RAF1 WT are the same as the four upper lines in Figure 1D. **D**: Senescent wild type WI38 cells were transfected with siVINK-1, siVINK-2, siCtle-1 and siCtle-2, whereas homozygously deleted (Δ/Δ #2) cells were transfected with siVINK-1 and siCtle-1. Total RNAs were prepared from two independent experiments and subjected to RNA-Seq. Differential gene expression (DGE) analysis was performed with the DESeq2 package. 1616 genes were found significantly deregulated by the two VINK siRNAs compared to the two controls in wild-type cells, and 2216 genes were significantly deregulated by the siVINK-1 siRNA in the Δ/Δ #2 cell line. These 2216 genes represent off-target genes since the siVINK-1 siRNAs does not decrease VINK expression in this cell line (see above). The Venn diagram shows the intersection between these two gene sets. Note that 720 out of the 1616 genes significantly deregulated by the VINK1 and VINK2 siRNAs in wild type cells are also significantly changed upon VINK-1 siRNA transfection in the deleted cell line.

Thus, removal of its target sequence abolished the effect of the VINK-1 siRNA on VINK expression, indicating that it represses the expression of its target RNA through base pairing to its target sequence. Of note, in the conditions we used to achieve VINK knock-down, VINK-1 siRNA induces prominent off-target effects with significant changes in the expression of thousands of genes upon siRNA transfection in the cell line in which the siRNA target site was deleted (Figure 4D).

### VINK siRNA requires cis-targeting to repress transcription of VINK

All together the above-described data unambiguously demonstrate that the VINK-1 siRNA represses VINK transcription by a process involving base-pairing. This suggests that VINK-1 siRNA targets VINK RNA and induces chromatin modifications at VINK promoter in cis that would lead to the repression of transcriptional initiation. However, we cannot formally rule out the possibility that the VINK-1 siRNA targets and induces the degradation by conventional RNAi mechanisms of a sub-population of VINK, an effect that would not be observed in our previous experiments because it only represents a minor proportion of VINK. If this population was able to indirectly regulate VINK expression, its depletion could then be signalized to the VINK promoter to repress VINK transcription. Of note, the VINK-1 siRNA targets a region located 300 bases upstream from a donor splice site (see Table S1 and S3), and may thus target a spliced product of VINK.

To formally demonstrate that VINK siRNAs act in cis, we raised by genome editing a cell line in which only one of the two alleles encoding VINK can be targeted by the VINK-1 siRNA, the other allele being deleted for the target sequence (that we called WI38-Δ/+). If the effects of siRNAs were due to base pairing at chromatin inducing local chromatin modifications, only the siRNA-targeted allele would be repressed. On the contrary, if the repression of VINK transcription was indirect, then the two alleles should be similarly regulated. We thus transfected WI38-Δ/+ cells with VINK-1 and VINK-2 siRNAs and performed RT-qPCR monitoring allele-specific VINK expression (see Figure 5A for the design of PCR primers). Each of the two siRNAs inhibited the expression of the wild type allele with a higher effect of the VINK-1 siRNA compared to the VINK-2 siRNA (Figure 5B), as in previous experiments (see Figure 1B-D). By contrast, on the deleted allele, while we still observed repression by the VINK-2 siRNA, repression by the VINK-1 siRNA was abolished (Figure 5C). The effect of the VINK-1 siRNA on total VINK expression in WI38- Δ/+ was weaker than on the wild type allele (compare Figure 5D with Figure 5B) or than in parental cells (compare Figure 5D with Figure 1B), as expected since only one allele was affected.

**Figure 5.**
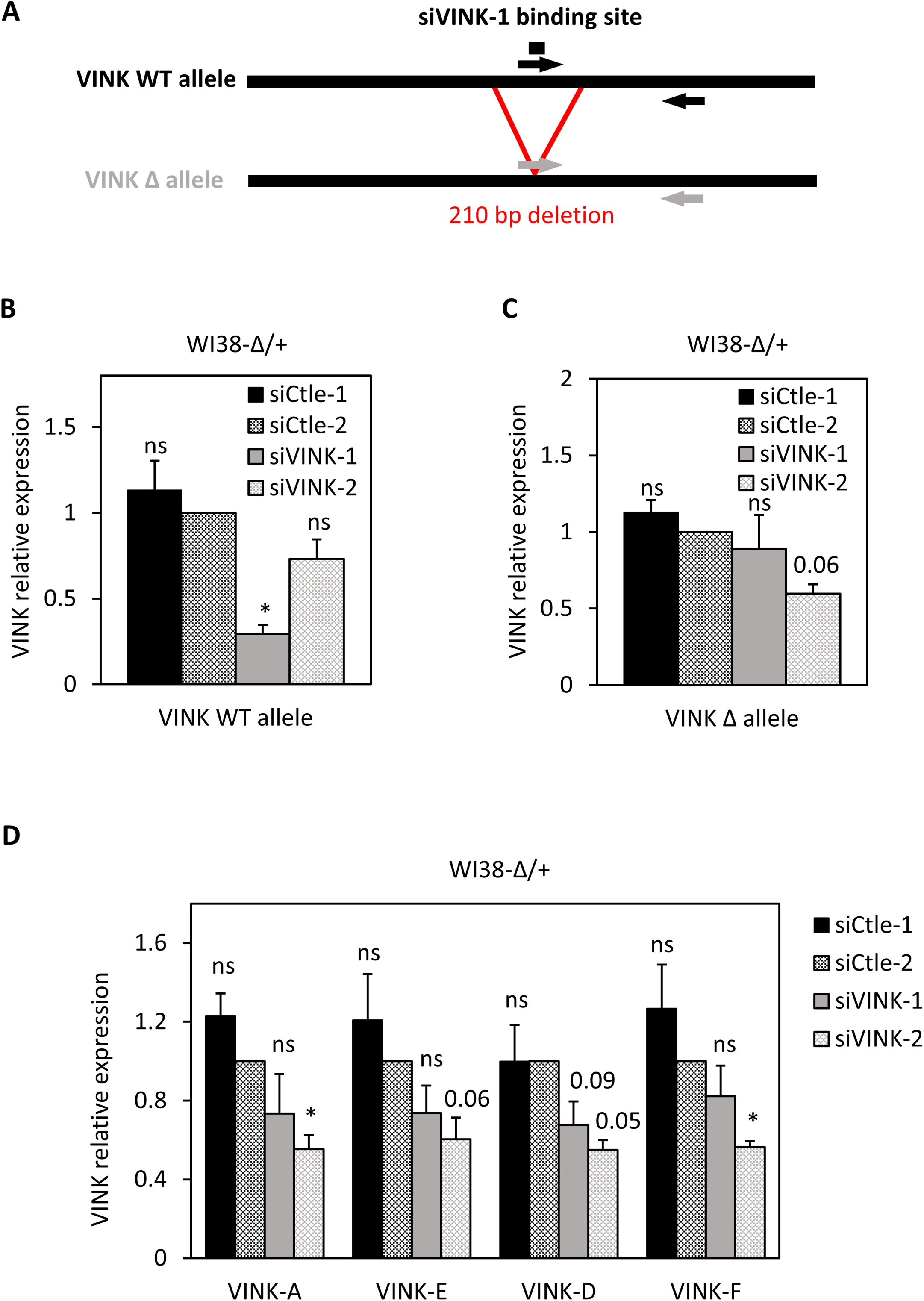
VINK siRNA requires a cis-targeting to chromatin to repress VINK transcription. **A**: Schematic representation of the two alleles of VINK present in WI38-Δ/+ with primers allowing their specific detection. The deleted allele is recognized by a primer located on the deletion junction. **B**: Senescent WI38-Δ/+cells were transfected with the indicated siRNAs. 72 hours later, total RNA was prepared and analysed by RT-qPCR for the expression of GAPDH mRNA or VINK wild type allele. The amount of VINK allele was standardised to GAPDH mRNA and calculated relative to 1 for cells transfected with the Ctle-2 siRNA. The means and standard deviations from 3 independent experiments are shown. *: p value < 0.05 compared to Ctle-2 siRNA ; ns: non-significant, p value > 0.1. **C**: Same as in B, except that the deleted VINK allele was analysed. The means and standard deviations from 3 independent experiments are shown. *: p value < 0.05 compared to Ctle-2 siRNA ; ns: non-significant, p value > 0.1; number: p value compared to Ctle2 siRNA. **D**: Same as in B and C, except that total VINK expression was analysed with the indicated primers. The means and standard deviations from 3 independent experiments are shown. *: p value < 0.05 compared to Ctle-2 siRNA ; ns: non-significant, p value > 0.1; number: p value compared to Ctle2 siRNA.

Altogether, these data thus demonstrate that the effect of VINK-1 siRNA on VINK transcription is mediated in *cis* by base-pairing of the siRNA to its target site on the chromatin-associated VINK RNA

### Ago1 and Ago2 are involved in siRNA-mediated VINK transcriptional repression

It was shown that transcriptional silencing by siRNA targeting promoters can involve either Ago1 or Ago2 (18, 31, 32). We thus investigated whether these proteins also participate in the transcription repression by siRNAs targeting VINK. We first tested whether Ago1 or Ago2 can be associated with chromatin in WI38 cells, since the nuclear localization of Ago proteins is somewhat debated. To that goal, we performed cell fractionation experiments to obtain cytoplasmic, nuclear soluble and chromatin protein fractions (Figure 6A). Cell fractionation was effective, since a-tubulin and GAPDH were mostly found in the cytoplasmic fraction, PARP and HDAC1/2 in the nuclear soluble fraction and histone H3 in the chromatin fraction. Interestingly, both Ago1 and Ago2 were detected in the three fractions, with an important amount present at chromatin. Comparison with the GAPDH or α-tubulin profiles indicates that this amount is unlikely due to contamination of the chromatin fraction with cytoplasmic proteins, and thus indicates that Ago1 and Ago2 are present at the chromatin in significant amounts. To test whether Ago1 or Ago2 expression was required for transcriptional repression by VINK1 siRNA, we made use of siRNAs targeting either Ago1 or Ago2. These two siRNAs were efficient and specific as observed by western blot (Figure 6B). We thus tested the effect of Ago1 and/or Ago2 knockdown on VINK repression by the VINK-1 siRNA. We found that siRNAs against Ago1 and Ago2 have a similar efficiency on their respective target whether or not they were transfected together with control or VINK-1 siRNAs, indicating that transfection of many siRNAs together did not affect their efficiency (Figure 6C and D). In contrast, the effect of the VINK-1 siRNA was weaker in the presence of either Ago1 or Ago2 siRNA (Figure 6C) and was nearly abolished when knocking-down both Ago1 and Ago2 (Figure 6D). Importantly, this last result was also observed when analyzing nascent RNA expression, confirming that it occurs at the transcriptional level (Figure 6E). Thus, these results indicate that both Ago1 and Ago2 are involved in siRNA-mediated transcriptional repression of VINK.

**Figure 6.**
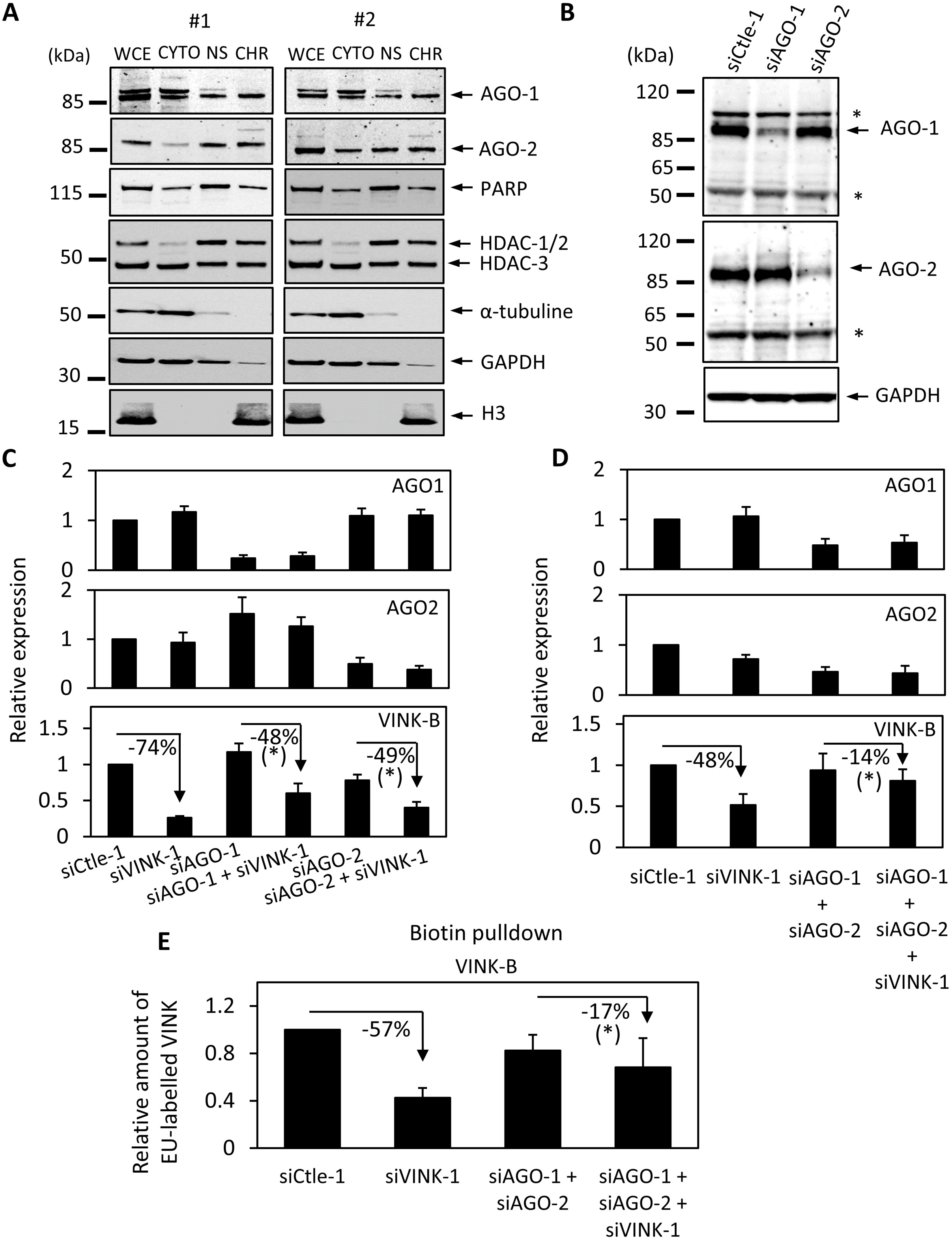
Ago1 and Ago2 are involved in VINK transcriptional repression by VINK siRNA. **A:** Senescent WI38 cells were recovered and subjected to a fractionation experiments. Whole cell extracts (WCE) and cytoplasmic (cyto), nuclear soluble (NS) and chromatin (CHR) fractions from the same amount of cells were analysed by western blot using the indicated antibodies. **B**: Senescent WI38 cells were transfected with 30 nM of the indicated siRNAs. 72 hours later, cells were harvested and total cell extracts were subjected to a western blot analysis with antibodies directed against Ago1, Ago2 or GAPDH as a loading control. The stars (*) indicate non-specific bands. Identical results were obtained in three independent experiments **C**: Senescent WI38 cells were transfected with 30 nM of the indicated siRNAs. The amount of siRNAs was kept constant at 60 nM using Ctle-1 siRNA. 72 hours later, total RNA was prepared and analysed by RT-qPCR for the expression of GAPDH, Ago1 or Ago2 mRNA or VINK. The amount of Ago1, Ago2 or VINK were standardised to GAPDH mRNA and calculated relative to 1 for cells transfected with the Ctle-1 siRNA. The means and standard deviations from 3 independent experiments are shown. *: p value < 0.05 when comparing the effect of siVINK-1 in cells depleted for Ago1 or Ago2 to its effect alone (paired student t test). **D**: Senescent WI38 cells were transfected with 25 nM of the indicated siRNAs. The amount of siRNAs was kept constant at 75 nM using Ctle-1 siRNA. 72 hours later, total RNA was prepared and analysed by RT-qPCR for the expression of GAPDH, Ago1 or Ago2 mRNA or VINK. The amount of Ago1, Ago2 mRNA or VINK were standardised to GAPDH mRNA and calculated relative to 1 for cells transfected with the Ctle-1 siRNA. The means and standard deviations from 6 independent experiments are shown. *: p value < 0.05 when comparing the effect of siVINK-1 in cells depleted for Ago1 and Ago2 to its effect alone (paired student t test). **E** Same as in D, except that nascent RNAs were recovered. The amount of VINK was standardised to GAPDH and calculated relative to 1 for cells transfected with the Ctle-1 siRNA. The means and standard deviations from three independent experiments are shown. *: p value < 0.05 when comparing the effect of siVINK-1 in cells depleted for Ago1 and Ago2 to its effect alone (paired student t test).

## Discussion

In this study, we show that siRNAs can induce efficient and specific transcriptional silencing of an RNA associated with chromatin. We observed the silencing of VINK promoter using siRNAs targeting the RNA in regions located thousands of bases downstream of the promoter, a phenomenom that, to our knowledge, has never been described so far in mammals. We further demonstrate the involvement of Ago1 and Ago2 in this process. This silencing was lost when the sequences targeted by the siRNA were deleted, ruling out off-target effects and demonstrating that base-pairing to its target sequence is required. Note that we cannot formally conclude whether this base pairing occurs on VINK RNA or on its transcribed template DNA.

The first question raised by our finding is whether the interference we have shown here with exogenously delivered siRNAs can also take place in natural conditions with endogenous siRNAs. Indeed, in various species, RNAi mechanisms have been shown to be involved in transcriptional silencing occurring at specific chromatin domains such as pericentric heterochromatin in fission yeast (4). Moreover, the involvement of endogenous antisense RNAs in the establishment or maintenance of heterochromatin silencing is well established (34, 39). Interestingly, it was recently proposed that tRNA-derived small RNAs (tsRNA) could be endogenous regulators of many genes, albeit by a different interference mechanism requiring cleavage by Ago2 (12), and other examples of endogenous siRNAs produced from protein-coding gene bodies have been described (40). Thus, we can speculate that endogenous RNAs complementary to chromatin-associated RNAs could be important regulators of the expression of these RNAs in mammals by triggering RNAi and transcriptional gene silencing.

To our knowledge, our study is the first to show that interfering RNAs targeting a RNA in regions far downstream of a promoter can induce can induce its transcriptional silencing in mammals. Such long-distance effects have been widely described in other species, in particular in plants. In these cases, it relies on RNA-dependent RNA polymerase (RdRP) which mediates amplification and propagation of double strand RNAs towards the 5’ end of RNAs (4). However, in mammals, RdRP is not conserved. What could thus be the mechanism involved in the process we describe here? It is known that double strand RNAs targeting the vicinity of a promoter (upstream promoter or the first exon) can lead to transcriptional silencing of the promoter through the induction of repressive epigenetic marks in a manner dependent on some RISC components (19, 22, 24). Here, we can speculate that the silencing of the promoter at distance of the siRNA target site could be achieved by chromatin folding that would bring the VINK/siRNA hybrid and the promoter in close proximity. As a consequence, this would change the chromatin landscape at the promoters, as proposed for miRNAs targeting the 3’ end of genes (26, 27). Of note, the siVINK-1 siRNA used in our studies targets a sequence close to an internal promoter, which is active in U2OS cells (data not shown) and probably also in WI38 (Ouvrard et al., in preparation). This second VINK TSS could favor its spatial proximity with the VINK promoter located more than 30 kb upstream. However, the fact that we observe transcriptional silencing with many different siRNAs argues against the importance of specific preexisting chromatin folding that would drive this silencing.

Contrary to the studies showing transcriptional gene silencing by miRNA targeting downstream regions of genes (26, 27), we observed a decrease in the presence of marks associated with transcription rather than an increase in repressive marks. Our data suggest the involvement of Ago proteins in this silencing. First, mere steric blocking by the VINK/siRNA hybrid is probably not sufficient for this silencing, since an antisense oligonucleotide targeting the same sequence as the VINK-1 siRNA does not induce transcriptional silencing, despite readily decreasing VINK expression (data not shown). Moreover, we found that Ago1 and Ago2 proteins depletion partially reversed VINK repression and that significant levels of Ago1 and Ago2 proteins are present at the chromatin in WI38 cells. However, by ChIP, we were not able to detect binding of Ago1 or Ago2 to the site targeted by the siRNA. We thus cannot formally rule out the possibility that the requirement of Ago proteins is indirect. Along this line, it would be interesting to analyse the relative importance of the siRNA seed domain versus other parts of the siRNA and whether 100% complementarity is required for silencing.

Despite the fact that intronic sequences are also found associated with chromatin, siRNAs directed against intronic sequences of pre-mRNAs are usually not effective to decrease the expression of pre-mRNAs (5, 6, 41). This finding first indicates that transcription of the target sequence is not sufficient to induce transcriptional silencing by siRNA, suggesting that the target of the siRNA is chromatin-associated RNA but not the DNA being transcribed. Splicing occurs co-transcriptionally leading to the degradation of introns, which thus greatly decrease the window of opportunity for the siRNA to bind efficiently its target RNA or to induce chromatin modifications when targeted. Moreover, siRNAs directed against a lncRNA did not induce transcriptional silencing, except when targeting the first exon (13). Thus, whatever the mechanism involved, it does not operate on any RNA associated with chromatin and we can speculate that the difference lies in the time of residence of the targeted RNA at chromatin.

The use of siRNAs to silence nuclear RNAs has been debated for long, because of the low levels of some RNAi proteins in the nucleus. As a consequence, siRNA-mediated degradation of RNA is mostly efficient in the cytoplasm. Nonetheless, as discussed above, siRNAs targeting promoters can in some instances repress transcription from the promoter they target. Here, we show that the promoter of chromatin-associated RNAs can be silenced using siRNAs targeting these RNAs in regions located far downstream of the promoter. Thus, siRNAs can represent interesting tools to investigate the function of chromatin-associated RNA by allowing their efficient and specific knock-down, even if the promoter is not well characterized or if the RNA is produced through multiple promoters. In addition, and coupled to a method allowing the degradation of RNA without interfering with any transcription step, it may help to distinguish between the role of the RNA itself and the role of its transcription, an ever-raising question in the field. Most importantly, since we found that siRNAs inhibit the transcription of the allele they target but not of the allele they do not target, they could allow studying *cis* effects of nuclear non coding RNAs or of their transcription, provided that an existing polymorphism or heterozygous genome editing, as we have done here, allows to target a single allele.

Note however that in the conditions we used to achieve the knock-down of VINK, VINK-1 siRNA induces prominent off-target effects with significant changes in the expression of thousands of genes upon siRNA transfection in the cell line in which the siRNA target site was deleted. Moreover, the use of two independent siRNAs was not sufficient to cope with off-target effects. Indeed, most of the genes showing expression changes with two independent siRNAs directed against VINK in wild type cells have a tendency to be changed in a similar way in the cell line deleted for the siRNA-1 target site (data not shown). These off-target effects are probably due to the fact that an siRNA can bind to and degrade target RNAs without perfect sequence complementarity, the seed region of only 6-7 nucleotides in the siRNA being involved in targeting numerous RNAs (42). It is possible that they are exacerbated when siRNAs do not have any target in the cytoplasm, such as for siRNAs directed against chromatin-associated RNAs or control siRNAs. Indeed, we also found thousands of DE genes between two different control siRNAs (data non shown). Importantly, we have not been able to dissociate the base pairing-dependent repression of VINK expression to off-target effects by changing the amount of siRNAs or the time after transfection at which we harvested cells (data not shown). It will thus be critically important for functional studies to design an experimental strategy ruling out off-target effects, such as overexpression of a siRNA-resistant form of the targeted RNA, or, as we performed here, removal of the siRNA target site by genome editing. When possible, this latter strategy is probably better since it maintains the normal production of the RNA at the normal place in the genome and in the nucleus, which is sometimes critical for the function of non-coding RNAs.

Finally, our findings suggest that siRNAs targeting chromatin-associated RNAs may have therapeutic potential. Indeed, the therapeutic value of siRNAs is currently widely investigated (43). Moreover, the relation of chromatin-associated RNAs with cancer progression has been documented in many instances, such as PVT1, H19, MALAT1 and HOTAIR (for a recent review see (44)). RNAs from the vlincRNA family are also overexpressed in cancer cells (45), and this overexpression may participate in allowing these cells to survive genotoxic stress (46). Targeting these RNAs with siRNAs may thus have therapeutic potential. Moreover, by modifying the chromatin landscape at their promoters and thus potentially reversing epigenetic information, it may have long-term benefits.

## Materials and Methods

### Cell culture

WI38-hTERT/ER-RAF1 immortalized fibroblastic cell line was kindly provided by C. Mann (47). They were grown in MEM supplemented with L-glutamine, non-essential amino acids, sodium pyruvate, penicillin-streptomycin and 10% of fetal bovine serum under 5% CO2 and 5% O2. Senescence was induced by treating cells with 20 nM of 4-hydroxy-tamoxifen for three days. U2OS cells (ATCC) were grown in DMEM containing Glutamax supplemented with sodium pyruvate, penicillin-streptomycin and 10% of foetal bovine serum under 5% CO2. WI38 cells and derivatives were transfected using the siRNA transfection reagent Dharmafect 4 (Dharmacon) following the manufacturer’s instructions with 100 nM of siRNA unless indicated in Figure legend and harvested 72 hours later. U2OS cells were transfected with the Interferin transfection reagent (Polyplus) or with Dharmafect 4 according to the manufacturers’ instructions with 50 nM of siRNA and harvested 48 hours later. For RNA stability analysis, actinomycin D (Sigma) was added at 10 µg/mL. siRNAs used are listed in Table S1:

### Cell fractionation, western blots and antibodies

Phosphatases and proteases inhibitors (thermofisher, # 78441) were added in all buffers. Fresh trypsinized cell were lysed in 5 cell volumes of lysis buffer (10 mM Tris pH8; 10 mM NaCl; 2 mM MgCl2). After 5 min on ice, 0.5% NP-40 was added. After another 10 min on ice, one third of the volume was kept as whole cell extract, the remaining lysate was centrifuged 10 min at 5000 rpm. The supernatant corresponded to the cytosolic fraction. The pellet (nuclei) was resuspended in 1 volume of buffer 3 (20 mM Hepes pH 7.9 ; 420 mM NaCl ; 1.5 mM MgCl2 ; glycerol 10%), then incubated 30 min on ice, shaking every 5 min. After 10 min of centrifugation at max speed, the supernatant was kept as nuclear soluble fraction. The pellet was the chromatin fraction. All the fractions (cytosol, nuclear soluble and chromatin) were equilibrated to the same final volume and same buffer compositions, adding 1% SDS in each. The whole cell extract was also adjusted to the same composition. The whole cell extract and chromatin were sonicated until viscosity disappeared. Laemmli sample buffer was added to the samples and the same volume of each fraction was loaded on an SDS-PAGE, except for the whole cell extract for which twice the volume of fractions was loaded.

Western blots were performed using standard protocols. Ago1 and Ago2 were detected using the monoclonal anti-AGO1 (6H1L4) antibody from Invitrogen (used at 1/1000) and the monoclonal anti-AGO2 (R.386.2) antibody from Invitrogen (1/1000), GAPDH was detected using the monoclonal anti-GAPDH (MAB374) antibody from Millipore (1/10000), PARP with anti-PARP from Cell signaling at 1/1000 (9542), HDAC1, HDAC2 and HDAC3 with anti-HDAC-3 from BD transduction lab at 1/500 (611125), H3 with anti-H3 from Abcam at 1/1000 (Ab1791) and α-tubulin with anti-tubulin from Sigma at 1/1000 (T6199).

For ChIP experiment, anti-RNA pol II was from Bethyl laboratories (A304-405A), anti-H3K27ac from Abcam (Ab4729), anti-H3K4me3 from Diagenode (pAb-003), anti-H3K9me3 from Abcam (Ab8898), anti-H3 from Abcam (Ab1791)and anti-H3K36me3 from Abcam (Ab9050). Purified rabbit IgG control was from Millipore (PP64B).

### Genome Editing

Genome editing of WI38 was performed with CRISPR-Cas9 tools in ribonucleic particle (RNP) format. Briefly, the Alt-R S.p. HiFi Cas9 Nuclease V3 (IDT) was assembled with equimolar amount of hybridized sgRNA (tracrRNA and crRNA from IDT) to generate RNPs. According to IDT recommendations, RNPs were transfected at 5 nM each with Lipofectamine CRISPRMAX (Thermofisher). Cells were harvested one week later to analyse the efficiency of genome editing. Genomic DNA was extracted with Master Pure purification kit (Epicentre) according to the manufacturer’s instructions, except that 250 µg of Proteinase K were used for cell lysis. Genome editing efficiency was checked by PCR on genomic DNA from the transfected population with the GoTaq G2 polymerase (Promega) using primers surrounding the 210 bp deleted region. PCR products were visualized on an agarose gel. Clones were then isolated and screened by PCR as described above. Two homozygous clones and one heterozygous clone were selected. The sequences of sgRNAs are the following ones:

sgRNA-1 : TAGGAGGCCAGTGCTCCAGG
sgRNA-2 : TTCCTGACTCTTGAAAACCA

### RNA extraction, reverse transcription and qPCR

Except when indicated, total RNA was prepared using the MasterPure RNA Purification Kit (Epicentre) according to the manufacturer’s instructions, except that 250 µg of Proteinase K was used for cell lysis. In Figure 2A, TRIzol (Sigma) was used. Briefly, adherent cells are directly lysed with 0.1 mL/cm^2^ of TRIzol during 10 min, the lysate is harvested before adding 0.1 mL of chloroforme per mL of TRIzol. After centrifugation, the upper phase is collected and the Epicentre kit is used for the following DNase digestion and subsequent precipitation. After total nucleic acids recovery, DNA was removed with a cocktail of DNaseI and Baseline Zero DNase supplemented with Riboguard RNase Inhibitor for 45 min at 37°C.

500 ng of total RNA were reverse transcribed with Superscript III reverse transcriptase (Invitrogen) according to manufacturer’s instructions. A condition without Superscript III was performed for each sample and analysed by qPCR using GAPDH exon 9 primers to check for DNA contamination. qPCR was performed in triplicate with TB green Premix Ex Taq from Takara and the Biorad CFX thermocycler. GADPH mRNA expression was used to normalize the results of RT-qPCR. Primers are listed in Table S2.

### Nascent RNA capture

Cells were incubated with 0.2 mM of ethynyl-uridine, a uridine analogue, for 1h. Cells were then harvested, total RNA was prepared and nascent RNA was purified from 2 µg of total RNA with the Click IT Nascent RNA capture kit from Life Technologies according to the manufacturer’s recommendations.

### Chromatin immunoprecipitation (ChIP)

ChIP was performed on 10.10^6^ transfected senescent WI38 cells as previously described (16), except that the nuclear lysis buffer was diluted twice before use, and sonicated nuclei were diluted 5 times in the dilution buffer. 10 million cells transfected with siRNA were crosslinked for 15 min using 1% formaldehyde directly in the culture medium. 0.125 M of glycine were then added for 5 min. After two washes with PBS, cells were scraped and frozen at −80°C. Cells were lysed with 3 ml of lysis buffer (5 mM Pipes pH 8, 85 mM KCl, 0.5% NP40) and homogenized 40 times with a dounce (20 times, pause 2 minutes, 20 times). After centrifugation, nuclear pellets were resuspended in 1.5 ml of nuclear lysis buffer (25 mM Tris pH 8.1, 5 mM EDTA, 0.5% SDS), and sonicated 10 times for 5s (power setting 0.5 ON 0.5 OFF and 50% amplitude, Branson Sonifier 250), to obtain DNA fragments of about 500 bp. DNA concentration was determined using a Nanodrop and samples were adjusted to the same concentration of chromatin. Samples were diluted five times in dilution buffer (0.01% SDS, 1.1% Triton X-100, 1.2 mM EDTA pH 8, 17 mM Tris pH 8.1, 167 mM NaCl) and precleared for 2 h with 250 μL of previously blocked 50% protein-A and protein-G beads (Sigma P-7786 and P-3296 respectively). Blocking was achieved by incubating the beads with 0.5 mg/ml of Ultrapure BSA and 0.2 mg/mL of salmon sperm DNA for 4 h at 4°C. 100 μL of chromatin were kept for inputs. 100 μg of pre-cleared samples per ChIP were incubated overnight for RNA pol II ChIP or 40 µg for histones ChIP with 4μg of antibody at 4°C. A mock sample without antibody was processed similarly. Then, 40 μL of blocked 50% A/G beads were added for 2h at 4°C to recover immune complexes. Beads were washed once in dialysis buffer (2 mM EDTA, 50 mM Tris pH 8, 0.2% Sarkosyl), five times in wash buffer (100 mM Tris pH 8.8, 500 mM LiCl, 1% NP40, 1% NaDoc) and twice in TE buffer (10 mM Tris pH 8, 1 mM EDTA). The bead/chromatin complexes were resuspended in 200 μL of TE buffer and incubated 30 min at 37 °C with 10 μg of RNase A (Abcam), as well as input DNA. Formaldehyde crosslink was reversed in the presence of 0.2% SDS at 70°C overnight with shaking. After 2 h of proteinase K (0.2 mg/ml) treatment at 45°C, immunoprecipitated and input DNA were purified on columns using Illustra GFX kit (GE Healthcare). All buffers for ChIP experiment were supplemented with EDTA-free protease inhibitor cocktail (Roche) and filtered 0.2 μM. Results were analysed by qPCR. Primers are listed in Table S2.

### Statistics

In all figures with statistical analyses, data were normalized to 1 relative to a control which was included in each experiment (usually a control siRNA). For significance analysis, the Log2 of fold change compared to this control was calculated in order to obtain data with a normal distribution. All experiments were then pooled together to calculate the variance of these experiments with a sufficient number of points. We then calculated using the student t test with this fixed variance the probability that the mean of each data set is equal to 0 (the value of the control), which represents the p value of the difference to the control. In some cases, we calculated using the paired student t test the probability that the mean of two populations is equal, which represents the p value of the difference between the two populations. A star is included in the Figure when the p value is below 0.05. The p value is indicated in the figure when it is between 0.05 and 0.1. When above 0.1, the difference is considered as non-significant and “ns” is indicated.

### RNA-seq

We performed strand-specific RNA-Seq method, relying on UTP incorporation in the second cDNA strand. For each sample, 5-10 μg of total RNA was submitted to EMBL-GeneCore, Heidelberg, Germany. Paired-end sequencing was performed by Illumina’s NextSeq 500 technology. Two replicates of each sample were sequenced.

The quality of each raw sequencing file (fastq) was verified with FastQC (48). Files were aligned to the reference human genome (hg38) in paired-end mode with STAR Version 2.5.2b and processed (sorting and indexing) with samtools (49). rDNA contamination was removed in the alignment process. Uniquely aligned reads were counted, per gene_id, using HT-seq Version 0.6.1 in a strand specific mode with the union count method parameter (50). The features’ annotation used for counts was constructed from NCBI refseq annotation gtf file from UCSC, taking the entire locus of reference for each gene. After removing lowly expressed counts (> 4 reads over the 4 samples), differential analysis was performed with DESeq2 Bioconductor R package, Version 1.22.1 (51). Counts were normalized using DESeq2 normalization method. Both treatment (siRNA VINK/siRNA Ctrl) and batch (replicates) effects were taken into account to estimate parameters and fit the model, in order to apply Wald test. Bigwig files were generated using rtracklayer Bioconductor-R package, Version 3.10, with 1 base pair bining and normalization based on the total number of read aligned. Spliced read of VINK were obtained by using GenomicAlignments (version 1.22.1) library in R script and human reference genome (BSgenome.Hsapiens.UCSC.hg38, version 1.4.1). Mapped paired-end reads were read from BAM with “readGAlignmentPairs” function. By using “njunc” and “junction” functions, the number of spliced reads and genomic coordinates of junctions were obtained (52). Spliced junctions were analysed as described in (15).

### Data availability

RNA-Seq data are available in the Geo database under the accession number # GSE197308

## Acknowledgements

This work was supported by a grant fom the Ligue Nationale Contre le Cancer to DT as an “équipe labellisée”. JO is supported by studentships from the French Ministry of Science and from the Fondation pour la Recherche Médicale (FRM). LM was supported by a fellowship from the Fondation de France. EN is a researcher at the INSERM.

The authors wished to thank Antonin Morillon, Hervé Seitz and all members of D Trouche’s lab for helpful discussions. We thank the genomic facility (BigA) from the CBI and Virginie Jouffret for RNA-Seq data analysis.

