## Supplemental Table 1 to 3 for "siRNAs targeting a chromatin-associated RNA induce its transcriptional silencing in human cells"

Supplementary Table S1 – List of siRNAs

| siRNAs name | Sequences | Position of 5' side of siVINK relative to estimated VINK TSS (nt) |
|---|---|---|
| siCtle-1 | S: GUCAGAGUAUCAUACGUAA-UU<br>AS: P-UUACGUAUGAUACUCUGAC-UU | |
| siCtle-2 | S: GACACAGUUCGAGUACUAA-UU<br>AS: P-UUAGUACUCGAACUGUGUC-UU | |
| siVINK-1 | S: CUUCUCAAGUUUCGAACAA-UU<br>AS: P-UUGUUCGAAACUUGAGAAG-UU | + 21 117 |
| siVINK-2 | S: GGGAAGGUAAGGAAAGAAA-UU<br>AS: P-UUUCUUUCCUUACCUUCCC-UU | + 163 298 |
| siARHGAP18-read_through | S: GGGCAGAGACUGAGACUUU-UU<br>AS: P-AAAGUCUCAGUCUCUGCCC-UU | |
| siAgo1 | S: CCCAGAUACUCCACUAUGA–UU<br>AS: P-UCAUAGUGGAGUAUCUGGG-UU | |
| siAgo2 | S: GGGUAAAGUUUACCAAAGA–UU<br>AS: P-UCUUUGGUAAACUUUACCC-UU | |

Supplementary Table S2 – List of primers

| Primers name | Sequences | Position of the amplicon relative to estimated VINK TSS (nt) |
|---|---|---|
| VINK-A | FW: CTGGACACACCCTAGGCAAG<br>RV: ATCCCTGTTGGTTATTCACAGC | + 207 to + 323 |
| VINK-B | FW: CCACAACCCCCATCTCTCTG<br>RV: CCCATCTTTGGTTCTTCTGTC | + 21 084 to + 21 390 |
| VINK-C | FW: TGGGTAGCATCTTCATCGGTG<br>RV: GAAGTGAGGGGAGTAGACAAG | + 22 497 to + 22 751 |
| VINK-D | FW: ACGCATGGGTGCCCTGTTTG<br>RV: ATAGATCTTGGAGGGTGCCAG | + 162 326 to + 162 445 |
| VINK-E | FW: ACCACTATACAATTGCAGCAGG<br>RV: CTGTCTGTTTAGGTACCTGGC | + 199 375 to + 199 539 |
| VINK-F | FW: GAGTGTCATCAGTAGTCTTAGCC<br>RV: CGAGGTCCAATATCCCTAACC | + 296 406 to + 296 540 |
| 420 bp encompassing the deletion product PCR primers | FW: GCATGAGTCATCTGGCTGTG<br>RV: CCCATCTTTGGTTCTTCTGTC | |
| WT VINK allele primers | FW: CCACAACCCCCATCTCTCTG<br>RV: CCCATCTTTGGTTCTTCTGTC | |
| Δ VINK allele primers | FW: ATATAGGAGGCCAGTGCTCCCCATG<br>RV: CCCATCTTTGGTTCTTCTGTC | |
| GAPDH exon 9 primers | FW: TGACAACGAATTTGGCTACAGC<br>RV: CTCTTCCTCTTGTGCTCTTGC | |
| GAPDH promoter primers | FW: AAATTGAGCCCGCAGCCTCC<br>RV: GCGACGCAAAAGAAGATGCG | |
| ARHGAP18-EX1 | FW: CAGGCGAGACAGGAACTTCTTT<br>RV: CTGCCTTTGCATGGCTGTTC | |
| ARHGAP18-INT1 | FW: GCTGCTGGAGTGAAATGTGG<br>RV: TGTTCAGCTAGTGAGAAGGTC | |
| ARHGAP18-read_through | FW: GTTCCTAGAGATTGATCTGAGG<br>RV: GAATAGACTTGGGTTGCCACG | |
| AGO1 | FW: CAGTGGACACCAACATCACC<br>RV: AAACGGTTGTCATCCCAAAG | |
| AGO2 | FW: CGCGTCCGAAGGCTGCTCTA<br>RV: TGGCTGTGCCTTGTAAAACGCT | |

| Junction | Donor splice site relative to estimated VINK TSS (nt) | Acceptor splice site relative to estimated VINK TSS (nt) | Number of reads containing this junction |
|---|---|---|---|
| 1 | + 330 | + 84479 | 78 |
| 2 | + 21 427 | + 84 479 | 16 |
| 3 | + 249 312 | + 250 728 | 16 |
| 4 | + 352 056 | + 352 076 | 15 |
| 5 | + 457 524 | + 465 489 | 36 |
| 6 | + 465 594 | + 506 188 | 34 |
| 7 | + 506 311 | + 550 054 | 21 |
| 8 | + 551 284 | + 603 617 | 10 |

Supplementary Table S3 : List of main spliced reads detected by RNA-Seq
The four siCtle samples from the RNA Seq shown in Figure 1D were combined and analysed for the detection of spliced reads. For each detected splicing junction, we calculated the number of reads which contained it. This table shows splicing junctions detected on more than 10 reads.
